# Domain-specific mutations in *unc-6/Netrin* differentially affect dorsal-ventral axon pathfinding in *Caenorhabditis elegans*

**DOI:** 10.64898/2026.07.14.738297

**Authors:** Kelsey M. Hooper, Scott G. Clark, Erik A. Lundquist

## Abstract

UNC-6/Netrin is a conserved regulator of dorsal-ventral axon and cell migrations. UNC-6 is composed of a Laminin N-terminal domain (LN), three epidermal growth factor repeats (EGF), and a Netrin C terminal domain (NC). Here, we identified missense mutations in distinct UNC-6 domains and assessed their roles in dorsal VD/DD motor axon guidance and ventral AVM axon guidance. A missense mutation in a conserved residue of the LN domain (G289D) resulted in dorsal and ventral axon guidance defects similar to *unc-6* null. A distinct missense mutation in the LN domain (S120F) was hypomorphic and strongly perturbed ventral AVM axon guidance with minimal effects on dorsal VD/DD axon guidance, showing that S120F is predominantly required for ventral guidance. Missense mutations altering conserved cysteine residues involved in disulfide bonding in the EGF domains were analyzed. EGF1(C321G) caused both ventral and dorsal axon guidance defects albeit weaker than *unc-6* null, indicating that EGF1 is required for both. EGF2(C347Y) strongly affected dorsal VD/DD axon guidance similar to *unc-6* null, with weaker perturbation of ventral AVM axon guidance. Previous results revealed that EGF3(C410Y) specifically disrupted dorsal axon guidance, a result that we confirmed. Our studies using missense mutations in the endogenous *unc-6* locus complement previous structure-function studies using transgenic expression, and identify domains specifically required for ventral AVM guidance (S120Y in the LN domain) and dorsal VD/DD axon guidance (C410Y in EGF3). The crystal structure of UNC-6 indicates conserved N-linked glycosylation at N114 and N128. Mutation of these sites in UNC-6 had no effect on dorsal ventral axon guidance, showing that they do not play a major role. However, the N114 and N128 mutations interacted genetically with *unc-40* and *unc-5* mutations, indicating that these glycosylation sites indeed have a role in UNC-6 signaling. Our results will inform studies on how these distinct UNC-6 domains interact with guidance receptors (*e.g.* UNC-40/DCC and UNC-5) and other extracellular molecules to mediate dorsal-ventral axon guidance.

## Introduction

UNC-6/Netrin and its receptors UNC-40/DCC and UNC-5 control dorsal-ventral axon guidance in *Caenorhabditis elegans* (HEDGECOCK *et al*. 1990; ISHII *et al*. 1992; SERAFINI *et al*. 1994; NORRIS AND LUNDQUIST 2011)(reviewed in (BOYER AND GUPTON 2018)). Classically, UNC-40/DCC receptor was thought to mediate ventral growth via attraction to UNC-6 (CHAN *et al*. 1996; KEINO-MASU 1996; KOLODZIEJ *et al*. 1996; DEINER *et al*. 1997), and the UNC-5 receptor was thought to control dorsal growth by repulsion from UNC-6 (LEUNG-HAGESTEIJN 1992; HAMELIN *et al*. 1993). Analysis of growth cone outgrowth in Netrin signaling mutants in *C. elegans* have revised this view and indicate that UNC-40/DCC and UNC-5 are both involved in ventral and dorsal guidance. In the Statistically-Oriented Asymmetric Localization (SOAL) model of growth toward Netrin, the protrusive activity of UNC-40/DCC is localized ventrally by UNC-6 in the HSN neuron, and UNC-5 focuses and maintains this polarity so that the HSN axon extends ventrally toward the UNC-6 source (KULKARNI *et al*. 2013; YANG *et al*. 2014; LIMERICK *et al*. 2017). In the Polarity/Protrusion model of dorsal growth away from UNC-6, Netrin first polarizes the VD growth cone through UNC-5 so that the pro-protrusive activity of UNC-40/DCC is localized dorsally, and then UNC-6 and UNC-5 maintain this polarity as the growth cone protrudes and grows dorsally away from the UNC-6/Netrin source (NORRIS AND LUNDQUIST 2011; NORRIS *et al*. 2014; GUJAR *et al*. 2018; MAHADIK AND LUNDQUIST 2023; HOOPER *et al*. 2025).

UNC-6/Netrin is similar to the N-terminus of Laminins and consists of a Laminin N-terminal (LN) domain, three Laminin-like Epidermal Growth Factor (EGF) repeats, and a Netrin-specific C-Terminal (NCD) domain (ISHII *et al*. 1992). Previous structure-function studies using domain deletions and mutations indicate that the LN domain and the EGF1 and EGF2 domains are required for both dorsal and ventral axon guidance, and that the EGF3 domain is required for required predominantly for dorsal axon guidance (HEDGECOCK *et al*. 1990; WADSWORTH AND HEDGECOCK 1996; LIM AND WADSWORTH 2002; NORRIS AND LUNDQUIST 2011). The NCD is not involved in dorsal or ventral guidance activity but rather acts to suppress ectopic branching (LIM *et al*. 1999; LIM AND WADSWORTH 2002; WANG AND WADSWORTH 2002). The UNC-5 extracellular Immunoglobulin like domain 2 (IG2) is predicted to physically interact with the UNC-6 LN and EGF1 domains, consistent with their roles in dorsal growth (PRIEST *et al*. 2024). Heparin binding at the IG1 and IG2 domains of UNC-5 stabilize the UNC-5-UNC-6 complex and exclude interaction with UNC-40 (PRIEST *et al*. 2024). Mammalian DCC interacts with Netrin1 at the LN, EGF2, and EGF3 domains (FINCI *et al*. 2014; XU *et al*. 2014), consistent with functional studies of *C. elegans* UNC-6 and UNC-40 (PRIEST *et al*. 2024).

Here, we report the identification, analysis, and sequencing of fourteen new *unc-6* alleles. Of these, four are missense mutations in specific UNC-6 domains. We characterized how these missense mutations affected dorsal VD/DD axon guidance and ventral AVM axon guidance. A glycine 289 to aspartic acid (G289D) substitution in the LN domain strongly disrupted both dorsal and ventral guidance, consistent with previous results (LIM AND WADSWORTH 2002). A serine 120 to phenylalanine (S120F) substitution in the LN domain was hypomorphic and predominantly perturbed ventral AVM axon guidance with minimal disruption of dorsal VD/DD axon guidance. Mutations of cysteines that make disulfide bonds in the EGF1(C321G) and EGF2(C347Y) disrupted both dorsal and ventral guidance, although to a lesser extent than the null *unc-6(ev400).* As reported previously (NORRIS AND LUNDQUIST 2011), the *unc-6(e78)* mutation that alters a disulfide-forming cysteine in the EGF3 domain (C410Y), strongly disrupted dorsal VD/DD guidance yet only partially disturpted AVM ventral guidance. In sum, our studies identified missense mutations that specifically affect the LN, EGF1, and EGF2 domains, and a novel S120F mutation in the LN domain that predominantly altered ventral AVM axon guidance. Our investigation using endogenous mutations complements previous structure-function studies with domain deletion rescuing transgenes (LIM AND WADSWORTH 2002).

The crystal structure of UNC-6 from protein produced in insect cells shows that UNC-6 is glycosylated at N114 and N128 (PRIEST *et al*. 2024). To test the role of N-glycosylation of UNC-6 in DV axon guidance, we changed N114 and N128 to alanine using Cas9 genome editing. We did not detect defects in VD/DD or AVM axon guidance in single and double mutants . However, we found via genetic interactions with *unc-40* and *unc-5* that UNC-6 function was reduced when these glycosylation sites were eliminated.

## Results

### Isolation and Analysis of *unc-6* alleles

We recovered fourteen new *unc-6* alleles in genetic screens for axon guidance mutants (see Materials and Methods). Each failed to complement the null *unc-6(ev400)* allele for the Uncoordinated (Unc) phenotype and were rescued for Unc by the *unc-6(+)* transgene *lqEx1241* (see Materials and Methods). We identified sequence changes in the *unc-6* gene in the mutants using Nanopore amplicon sequencing (see Materials and Methods). *zd17, zd116, zd146,* and *zd154* contain missense mutations (Table 1 and Figure 1) The mutation in *zd17* changed serine 120 to phenylalanine and the mutation in *zd146* changed glycine 289 to aspartic acid, both in the laminin N-terminal (LN) domain. Both sites are conserved in Netrin and Laminin sequences across phylogeny including vertebrates (SHAW *et al*. 2021). Indeed, the equivalent of UNC-6 glycine 289 is affected by pathogenic missense mutations in human Laminin α2 (limb-girdle-type dystrophy) (GAVASSINI *et al*. 2011). Mutation of Drosophila Laminin β1 at this position led to cardiac development defects (HOLLFELDER *et al*. 2014).

**Figure 1.**
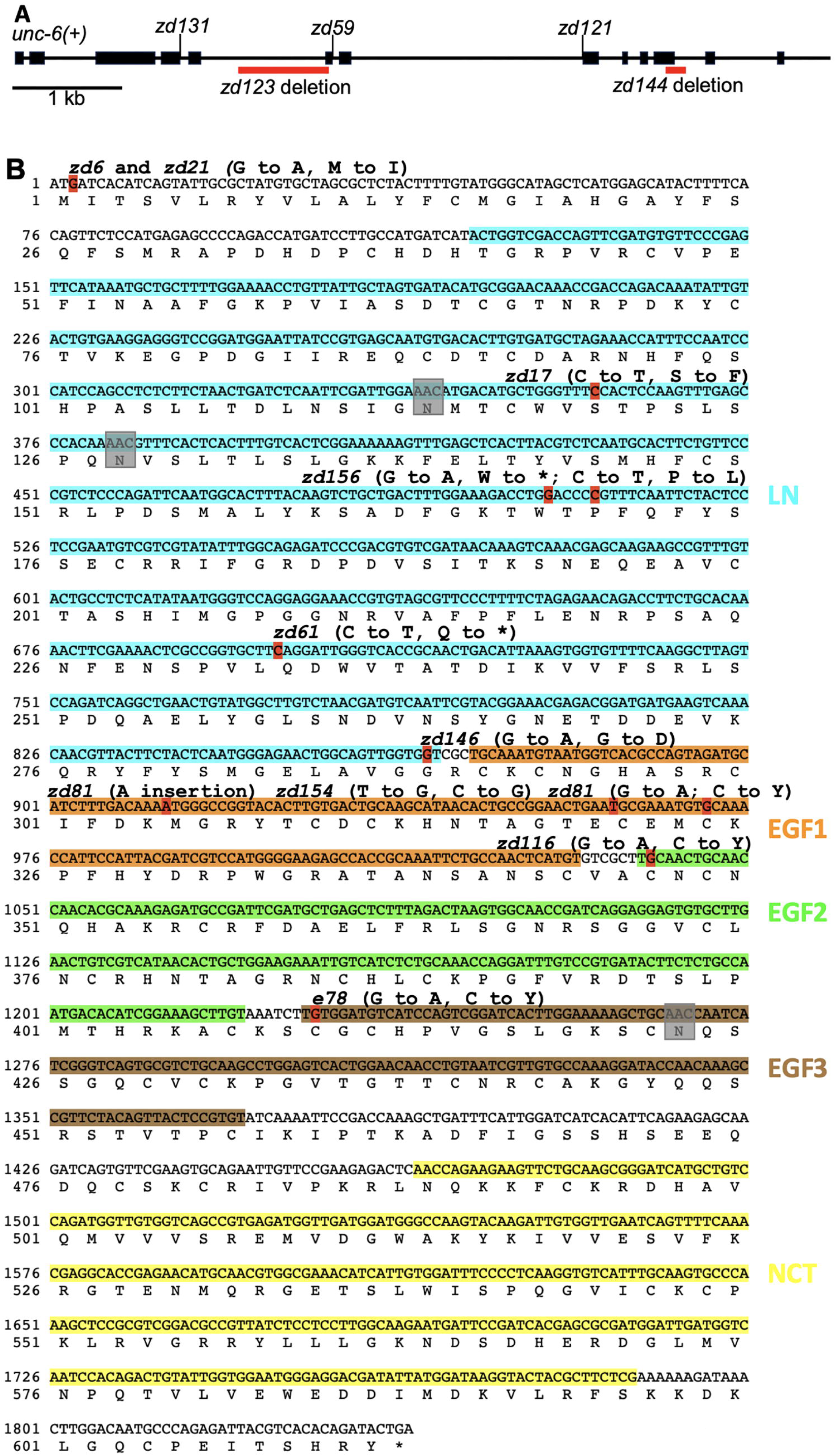
The *unc-6* locus and new *unc-6* alleles. A) A diagram of the *unc-6* locus. Exons are represented by boxes, and introns by lines. The *zd123* and *zd144* deletions are shown, and the *unc-6* mutations affecting splice sites are indicated. B) The *unc-6* cDNA sequence. The Laminin N terminal domain (LN) is highlighted in blue; the first EGF domain (EGF1) in orange; EGF2 in green; EGF3 in brown; and the Netrin C-terminal domain (NCT) in yellow. The locations of new *unc-6* alleles are shown above the sequence and indicated in red. The three conserved potential N-glycosylation sites are highlighted by grey boxes.

**Table 1.**
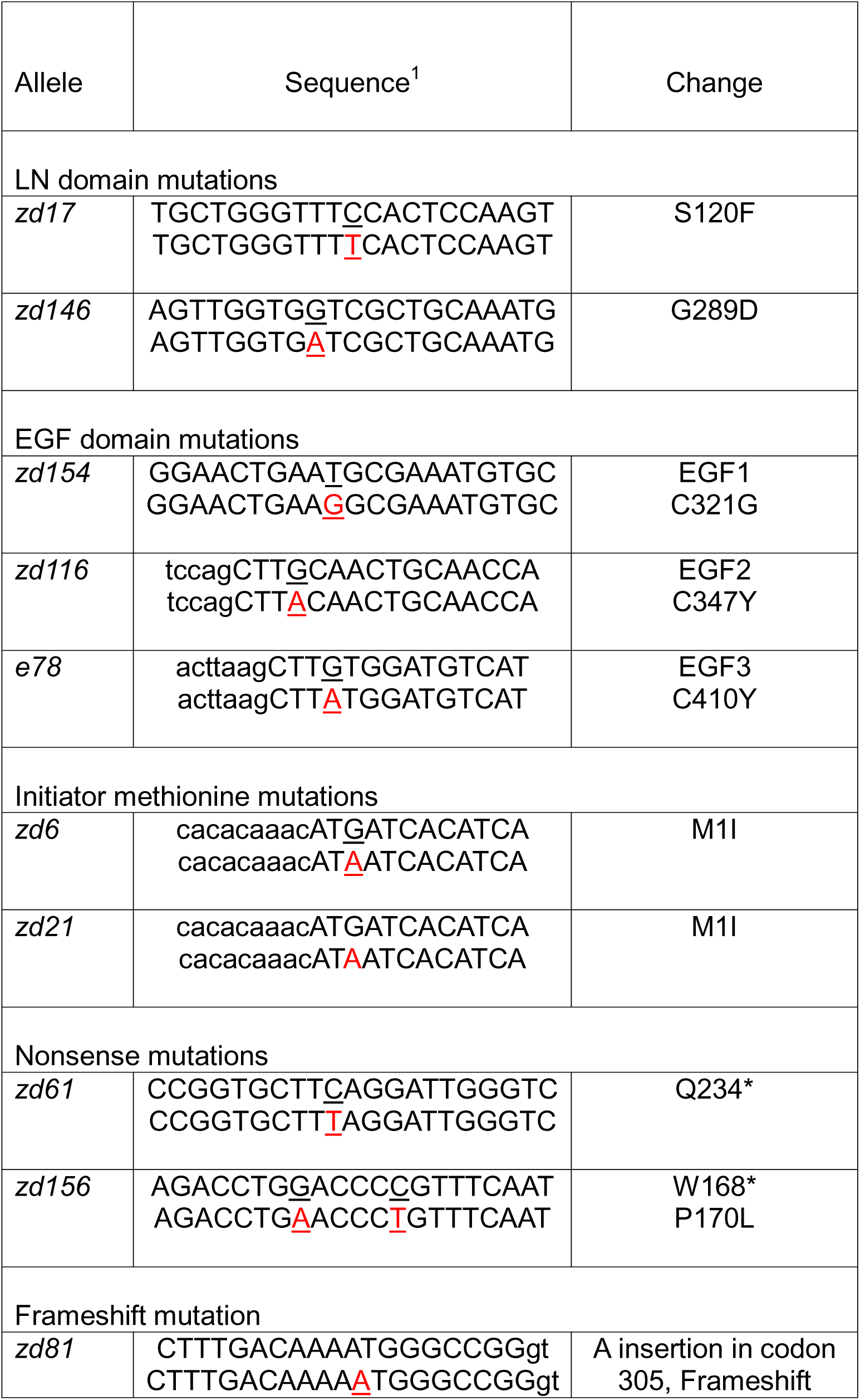
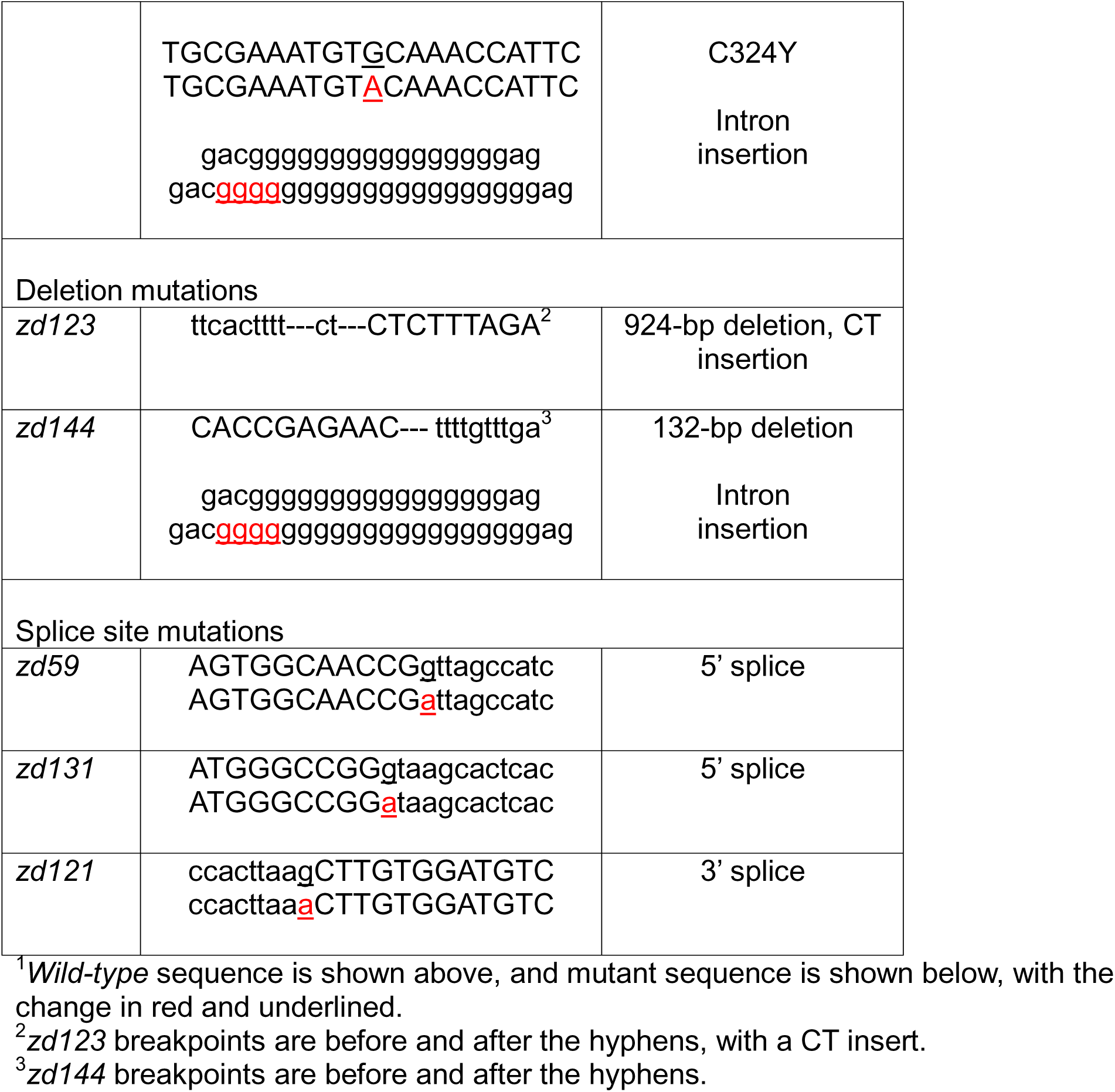
Fourteen new *unc-6* alleles.

*zd154* changed cysteine 321 to glycine in the first EGF domain, and *zd116* changed cysteine 347 to tyrosine in the second EGF domain. The *unc-6(e78)* allele was previously shown to change cysteine 410 to tyrosine in the third EGF domain. The cysteines affected by *zd154* (EGF1), *zd116* (EGF2), and *e78* (EGF3) are each involved in disulfide bonds that contribute to the structure of the EGF domains (Figure 2A-C).

**Figure 2.**
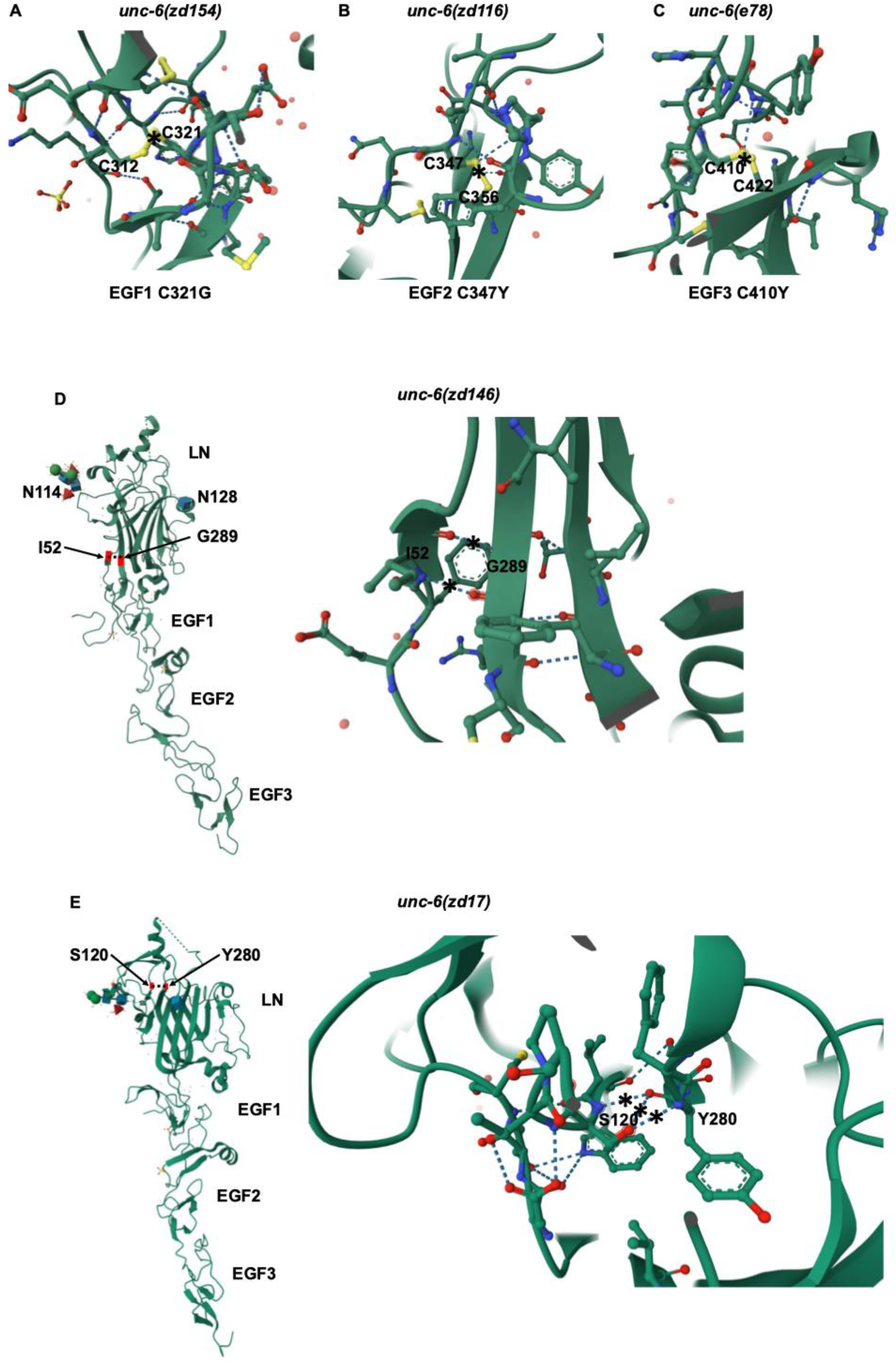
Crystal structures of UNC-6 regions affected by new *unc-6* missense mutations. Images were generated by Mol* from the X-ray crystal structure of the UNC-6 LN and EGF domains in the Protein Data Bank (BERMAN *et al*. 2000; PRIEST *et al*. 2024) (pdb00008EDK). A) EGF1 residue Cysteine 321, changed to a glycine in *unc-6(zd154)*, normally makes a disulfide bond with cysteine 312 (asterisk). B) EGF2 residue cysteine 347, changed to a tyrosine in *unc-6(zd116)*, normally makes a disulfide bond with cysteine 356 (asterisk). C) EGF3 residue cysteine 410, changed to a tyrosine in *unc-6(e78)*, normally makes a disulfide bond with cysteine 422 (asterisk). D) In the LN domain, glycine 289, changed to aspartic acid in *unc-6(zd146)*, makes 2 hydrogen bonds with isoleucine 52 in an adjacent β strand (dotted line and asterisks). The potentially N-glycosylated residues asparagine 114 and asparagine 128 in the LN domain are also indicated. E) Serine 120 in the LN domain, changed to phenylalanine in *unc-6(zd17)*, makes three hydrogen bonds with tyrosine 380 in an adjacent β strand in the “jellyroll” β-barrel structure.

The remaining ten *unc-6* alleles harbor mutations that are predicted to produce non-functional UNC-6 protein because of failure to initiate translation, premature termination of translation, or aberrant mRNA splicing. *zd6* and *zd21* contain the same missense mutation that converts the initiator methionine to isoleucine (1MI) (Table 1 and Figure 1). There is an in-frame ATG at position 16 which, if used, would result in a molecule lacking a signal peptide. *zd61* and *zd156* contained nonsense mutations that introduce premature stop codons (Table 1 and Figure 1). *zd156* had a second C to T mutation resulting in a P170L missense mutation five bases downstream of the nonsense mutation (Table 1). *zd81* contained an A residue insertion in codon 305 resulting in a frameshift and in-frame premature stop codon six codons downstream. *zd81* also contained a missense mutation (C324Y) downstream of the frameshift and an insertion of 4 G residues in intron 7. *zd123* contained a 924-bp deletion with a 2-bp insertion that removed 47 bp of coding sequence in addition to intron sequence (Table 1 and Figure 1). *zd144* contained a 132-bp deletion that removed 94 bases of coding region as well as intron sequence (Table 1 and Figure 1). *zd144* also contained the insertion of four G residues in intron 7 identical to *zd81.* zd59, zd121, and zd131 contained point mutations predicted to disrupt splicing of unc-6 mRNA: zd59 and zd131 altered an invariant G in the 5’ splice donor sequence and zd121 changed an invariant G in the 3’ splice accepter sequence (Figure 1 and Table 1).

### Dorsal-ventral axon guidance defects in LN domain *unc-6* mutants

The axons of the VD and DD motor neurons are a model for dorsal growth cone migration away from the UNC-6 source in the ventral nerve cord. The 19 VD and DD motor neurons extend axons dorsally from the ventral nerve cord to the dorsal nerve cord. In wild type, we observed an average of 16.21 commissures due to fasciculation of some axons into single commissures (Table 2 and Figure 3A). Null *unc-6(ev400)* mutants displayed severe defects in VD/DD dorsal axon guidance, with very few axons reaching the dorsal nerve cord (an average of 0.02 per animal, or only two observed in 100 animals), and an average of only 6.22 per animal showing any extension dorsally from the ventral nerve cord (Table 2 and Figure 3B).

**Figure 3.**
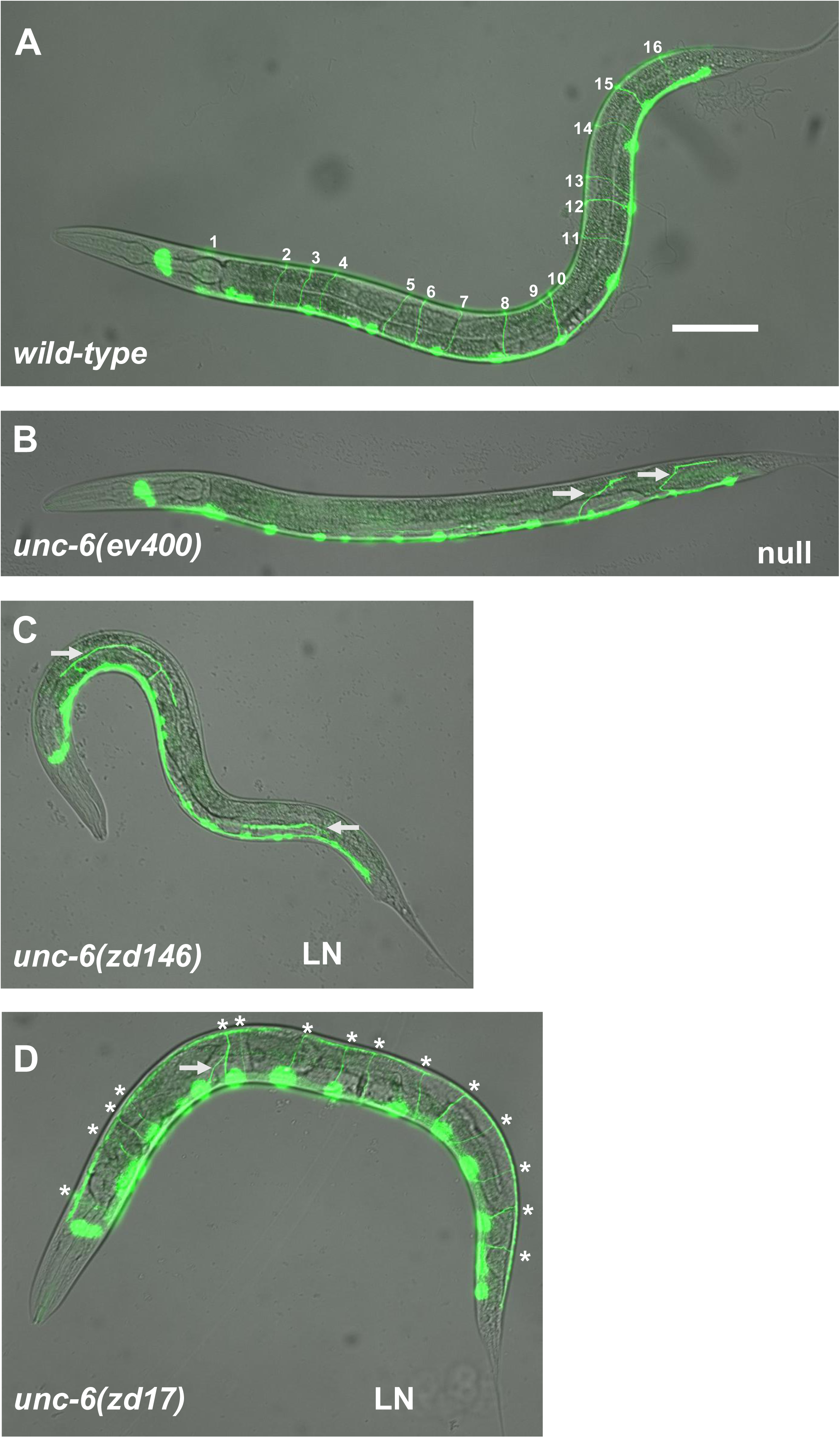
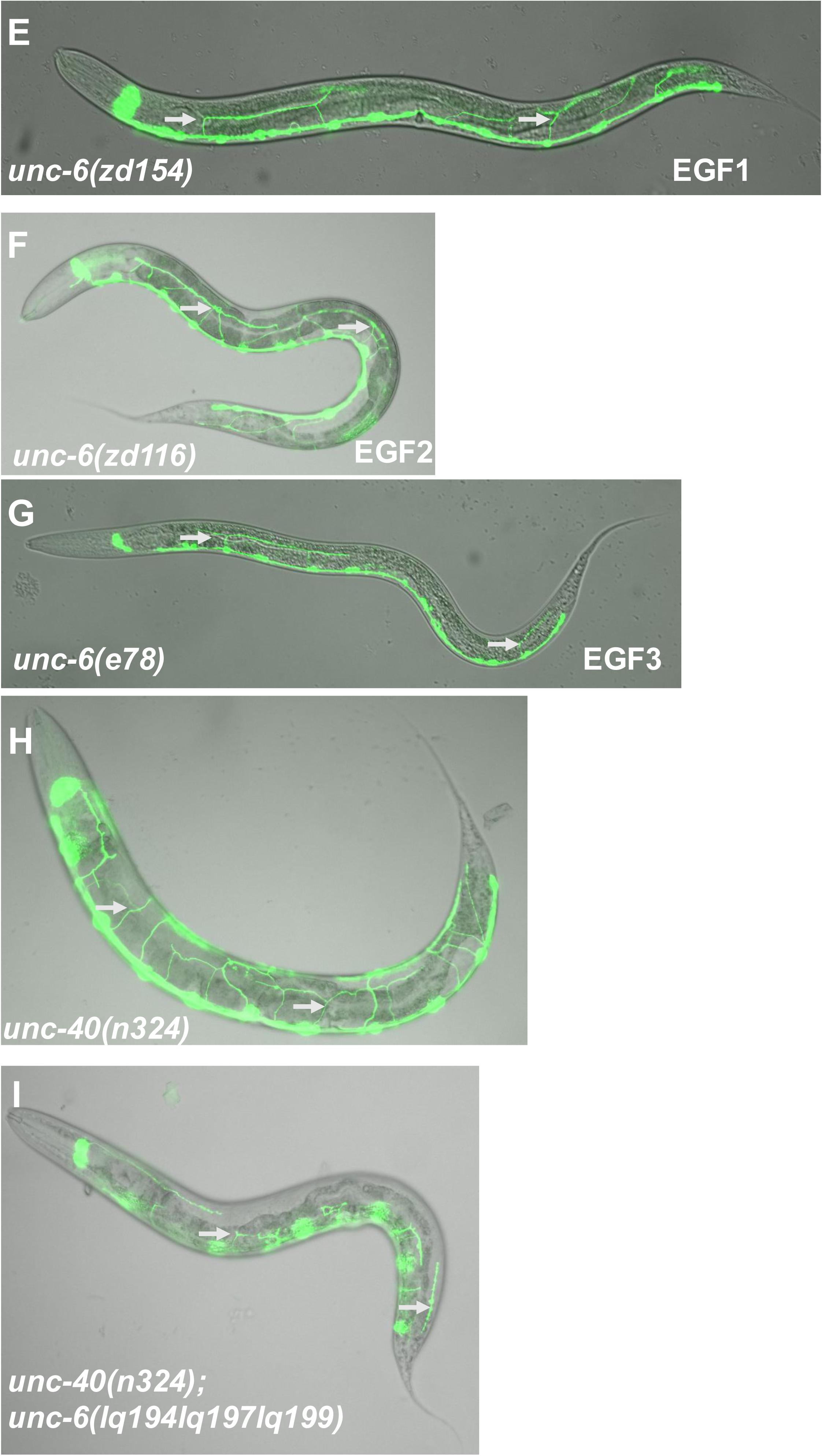
VD/DD dorsal axon guidance defects in *unc-6* mutants. Micrographs of the *juIs76[Punc-25::gfp]* transgene expressed in the GABA-ergic motoroneurons with a merged differential interference contrast micrograph are shown. Anterior is to the left and dorsal up. The scale bar in A indicates 50μm for all micrographs. The domain affected by each mutant is indicated. A) A wild-type animal displaying 16 visible VD/DD commissural tracts from the ventral nerve cord (VNC) to the dorsal nerve cord (DNC) (numbers). B) An *unc-6(ev400)* null mutant. Arrows point to axons that emerged from the VNC, but none reached the DNC. C) An *unc-6(zd146)* mutant with axons emerging from the VNC (arrows) but none reaching the DNC. D) An *unc-6(zd17)* mutant showed 15 normal VD/DD axon commissures (asterisks) with one misguided axon (arrow). E) An *unc-6(zd154)* mutant with some misguided axons emanating from the VNC (arrows). F) An *unc-6(zd116)* mutant with many axons emanating from the VNC but not reaching the DNC (arrows). G) An *unc-6(e78)* mutant with some axons emanating from the VNC (arrows). H) An *unc-40(n324)* mutant with defective VD/DD axon guidance (arrows), many of which extend away from the VNC and reach the DNC. I) An *unc-40(n324); unc-6(lq194lq197lq199)* mutant with severe VD/DD axon guidance defects (arrows), with few axons extending away from the VNC.

**Table 2.** VD/DD and AVM axon guidance defects.

| Genotype | Average # VD/DD emerging | Average # VD/DD reaching | % defective AVM ventral guidance |
| --- | --- | --- | --- |
| *wild-type* | 16.02 | 15.95 | 0 |
| *unc-6(ev400)*<br>Null | 6.22 | 0.02 | 72 |
| *unc-6(zd146)*<br>LN | 5.57 | 0.04 | 67 |
| *unc-6(zd17)*<br>LN | 15.82 | 9.94 | 30 |
| *unc-6(zd154)*<br>EGF1 | 6.05 | 0.02 | 35 |
| *unc-6(zd116)*<br>EGF2 | 12.80 | 1.78 | 18 |
| *unc-6(e78)*<br>EGF3 | 4.54 | 0.03 | 8 |
| *unc-6(zd144)*<br>C-terminal deletion | 8.41 | 0.04 | 45 |
| *unc-6(lq194) N114A* | 15.98 | 15.92 | 0 |
| *unc-6(lq196) N128A* | 16.00 | 15.91 | 0 |
| *unc-6(lq200) N423A* | 15.87 | 15.78 | 0 |
| *unc-6(lq194lq197) N114A, N128A* | 16.03 | 15.91 | 0 |
| *unc-6(lq194lq197lq199) N114A, N128A, N423A* | 15.76 | 15.58 | 0 |
| *unc-40(n324)* | 15.76 | 9.88 | 11 |
| *unc-40(n324); unc-6(N114A)* | 15.64 | 7.13 | ND |
| *unc-40(n324); unc-6(N128A)* | 15.61 | 7.23 | ND |
| *unc-40(n324); unc-6(N423A)* | 15.37 | 4.66 | ND |
| *unc-40(n324); unc-6(N114A N128A)* | 15.70 | 6.41 | ND |
| *unc-40(n324); unc-6(N114A N128A N423A)* | 15.13 | 3.60 | 17 |
| *unc-5(ev480)* | 10.72 | 6.61 | 0 |
| *unc-5(ev480); unc-6(N114A)* | 13.34 | 4.06 | 2 |
| *unc-5(ev480); unc-6(N128A)* | 14.04 | 4.68 | 1 |
| *unc-5(ev480); unc-6(N423A)* | 13.57 | 4.82 | 1 |
| *unc-5(ev480); unc-6(N114A N128A)* | 13.48 | 4.54 | 3 |
| *unc-5(ev480); unc-6(N114A N128A N423A)* | 9.76 | 2.82 | 5 |
| *unc-5(e791)* | 0.25 | 0 | 4 |
| *unc-5(e791); unc-6(N114A N128A N423A)* | ND | ND | 7 |
| Statistical comparisons | ANOVA | ANOVA | Fisher's Exact |
| zd146 vs. ev400 | NS | NS | NS |
| zd17 vs. ev400 | $p < 0.0001$ | $p < 0.0001$ | $p < 0.0001$ |
| zd154 vs. ev400 | NS | NS | $p < 0.0001$ |
| zd116 vs. ev400 | $p < 0.0001$ | $p < 0.0001$ | $p < 0.0001$ |
| zd144 vs. ev400 | $p < 0.0001$ | NS | $p = 0.0002$ |
| zd116 vs. zd154 | $p < 0.0001$ | $p < 0.0001$ | $p = 0.0100$ |
| zd116 vs. e78 | $p < 0.0001$ | $p < 0.0001$ | $p = 0.0568$ |
| e78 vs. ev400 | $p < 0.0001$ | NS | $p < 0.0001$ |
| e78 vs. zd17 | $p < 0.0001$ | $p < 0.0001$ | $p < 0.0001$ |
| lq194 vs. wild-type | NS | NS | NS |
| lq196 vs. wild-type | NS | NS | NS |
| lq200 vs. wild-type | NS | NS | NS |
| lq194lq197 vs. wild-type | NS | NS | NS |
| lq194lq197lq199 vs. wild-type | NS | $p = 0.0261$ | NS |
| n324; lq194 vs. n324 | NS | $p < 0.0001$ | ND |
| n324; lq196 vs. n324 | NS | $p < 0.0001$ | ND |
| n324; lq200 vs. n324 | NS | $p < 0.0001$ | ND |
| n324; lq194lq197 vs. n324 | NS | $p < 0.0001$ | ND |
| n324; lq194lq197lq199 vs. n324 | NS | $p < 0.0001$ | NS |
| ev480; lq194 vs. ev480 | $p < 0.0001$ | $p < 0.0001$ | NS |
| ev480; lq196 vs. ev480 | $p < 0.0001$ | $p < 0.0001$ | NS |
| ev480; lq200 vs. ev480 | $p < 0.0001$ | $p < 0.0001$ | NS |
| ev480; lq194lq197 vs. ev480 | $p < 0.0001$ | $p < 0.0001$ | NSq |
| ev480; lq194lq197lq199 vs. ev480 | $p = 0.0037$ | $p < 0.0001$ | $p = 0.0594$ |
| e791 vs. e791; lq194lq197lq199 | ND | ND | NS |

The axon of the AVM mechanosensory neuron is a model for ventral axon growth toward the UNC-6 source in the ventral nerve cord. The AVM cell body resides laterally, and extends an axon ventrally to ventral nerve cord (Figure 4A). In *unc-6(ev400)* null mutants, the AVM axon failed to extend ventrally in 72% of animals and instead extended laterally toward the anterior (Table 2 and Figure 4B).

**Figure 4.**
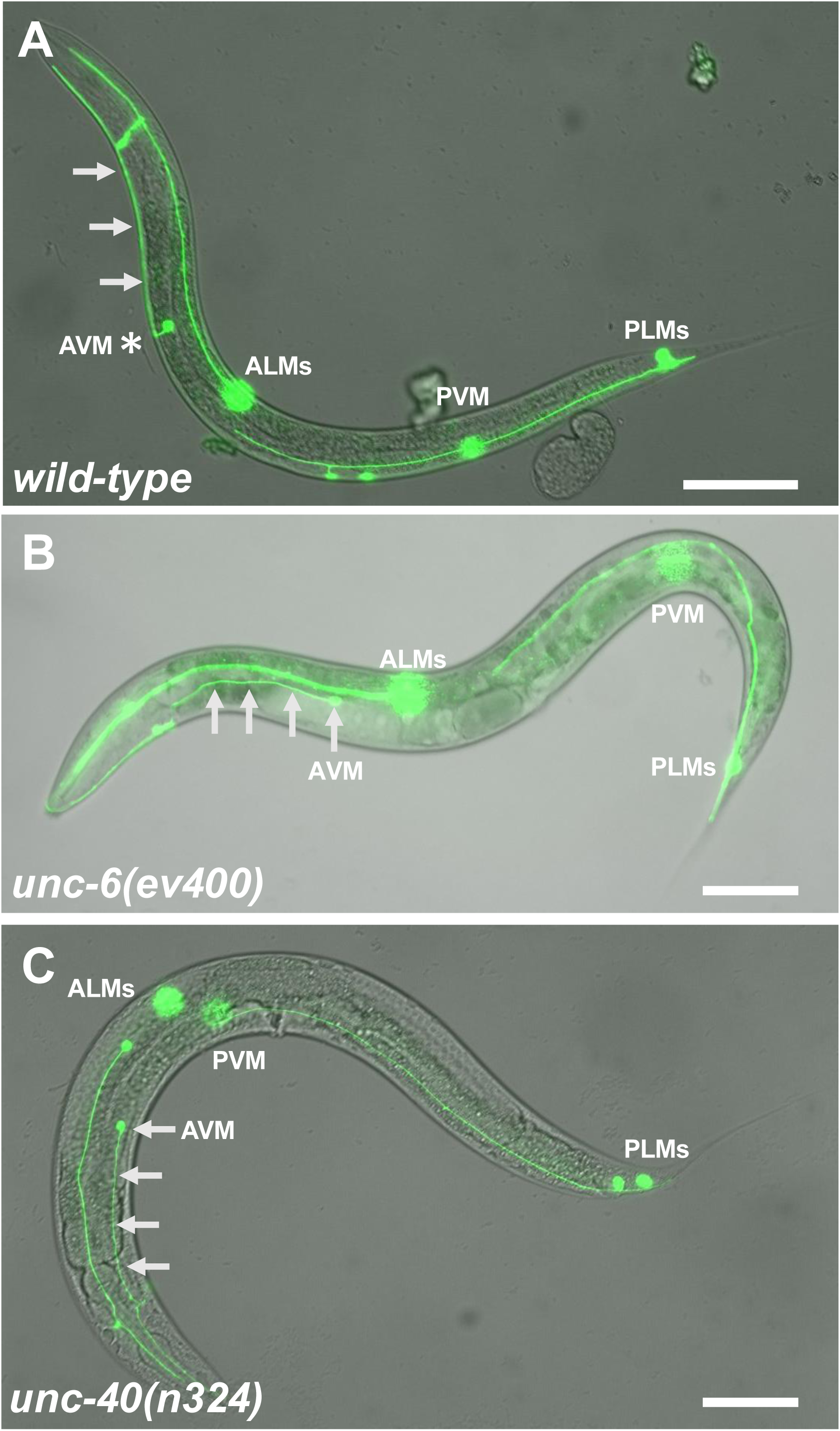
AVM ventral axon guidance defects. Merged GFP fluorescent and DIC micrographs are shown as described in Figure 3. The scale bars represent 50μm. A and B are the *zdIs5[Pmec-4::gfp]* transgene and C is the *zdIs4[Pmec-4::gfp] zdIs4[Pmec-4::gfp]* transgene, both expressed in the mechanosensory neurons. The positions of the AVM cell body is indicated, as are the ALMs, PLMs, and PVM, sometimes out of focus. Anterior is left and dorsal is up. A) A *wild-type* animal in which the AVM axon extends directly ventrally from the cell body (asterisk). The AVM axon then runs anteriorly in the VNC (arrows). B) An *unc-6(ev400)* animal in which the AVM axon fails to extend ventrally and instead extend anteriorly out of the AVM cell body along the lateral body wall (arrows). B) An *unc-40(n324)* animal in which an AVM axon fails in ventral extension and instead extends anteriorly along the lateral body wall (arrows). The PVM cell body is also anteriorly displaced due to the previously-described defective migration of the QL neuroblast and descendants in *unc-40* mutants.

We scored axon defects in *unc-6(zd17)* and *unc-6(zd146)*, missense mutations in the LN domain. *unc-6(zd146)* animals were strongly uncoordinated, and displayed severe defects in VD/DD dorsal axon outgrowth not significantly different from the *unc-6(ev400)* null (Table 2 and Figure 3C). Likewise, *unc-6(zd146)* displayed strong AVM ventral axon guidance defects not significantly different from the *unc-6(ev400)* null (Table 2). These results indicate that *unc-6(zd146)*, a G289D missense mutation in the LN domain, is a strong loss of function mutation, comparable to a null. In the UNC-6 crystal structure, G289 makes two hydrogen bonds with I52 in a neighboring β-strand near the base of the “jelly roll” β-barrel core of the LN domain (Figure 2D). While the peptide backbone of the G residue is involved in the hydrogen bonding, insertion of a larger D residue might disrupt this interaction.

*unc-6(zd17)* mutants had less severe uncoordination compared to *unc-6(ev400)* or *unc-6(zd146)* mutants. VD/DD dorsal guidance was only mildly affected in *unc-6(zd17)*, with an average of 15.82 axons emanating from the ventral nerve cord and an average of 9.94 reaching the dorsal nerve cord, significantly less severe than the *unc-6(ev400)* null (Table 2 and Figure 3D). However, AVM ventral guidance failed in 30% of *unc-6(zd17)* mutants (Table 2). Our data shows that *unc-6(zd17)* strongly perturbed ventral axon guidance yet only weakly perturbed dorsal axon guidance.

The *zd17* mutation altered S120, a residue conserved in the LN domains of Netrins and laminins. In the UNC-6 crystal structure, S120 forms three hydrogen bonds with Y280 in an adjacent β-strand, between the β-barrel “jelly roll” core of the LN domain and a nearby domain that is potentially subject to glycosylation (Figure 2E). Two of the three hydrogen bonds involve the serine side group. Insertion of a larger tyrosine residue likely disrupts these interactions.

### Dorsal-ventral axon guidance defects in EGF domain mutants

The cysteines affected by *unc-6(zd154), unc-6(zd116),* and *unc-6(e78)* are each involved in disulfide bonds with cysteines in the core structure of the three EGF domains (Figure 2A-C). These mutations might predominantly affect the EGF domain in which they reside. Previous phenotypic analysis indicate that *unc-6(e78)* strongly affected dorsal axon guidance, with weaker effects on ventral axon guidance. For example, very few VD/DD commissures reached the dorsal nerve cord (only three in 100 animals scored), not significantly different from the *unc-6(ev400)* null, and only 4.54 on average emanated from the ventral nerve cord, significantly fewer than the *unc-6(ev400)* null (Table 2 and Figure 3G). However, only 8% of AVM axons failed to extend ventrally, significantly fewer than the 72% for *unc-6(ev400)* and the 30% for *unc-6(zd17)*) (Table 2). These results show that the EGF3 domain is required primarily for dorsal growth away from UNC-6, and support the idea that the *unc-6(zd17)* mutation might primarily affect ventral growth.

*unc-6(zd154)* affected a conserved cysteine in the first EGF domain. *unc-6(zd154)* affected both dorsal and ventral growth. *unc-6(zd154)* displayed on average 6.02 VD/DD axons emanating from the ventral nerve cord, similar to the *unc-6(ev400)* null, and did not differ significantly from the *unc-6(ev400)* null for the number of axons reaching the DNC (Table 2 and Figure 3E). *unc-6(zd154)* had significantly weaker effects on AVM ventral guidance compared to the *unc-6(ev400)* null, with 35% of AVM axons that failed to extend ventrally in *unc-6(zd154)* (Table 2).

*unc-6(zd116)* altered a conserved cysteine in the second EGF domain, the analogous cysteine affected by *unc-6(e78)* in the third EGF domain. *unc-6(zd116)* affected both dorsal and ventral growth but was significantly weaker than the *unc-6(ev400)* null and the *unc-6(zd154)* EGF1 mutant (Table 2 and Figure 3G). Furthermore, 18% of AVM axons failed to grow ventrally in *unc-6(zd116)* (Table 2), significantly weaker than the *unc-6(ev400)*. These data indicate that EGF domains 1 and 2 are required for both dorsal and ventral axon guidance, although perturbation of EGF1 more strongly affected dorsal guidance than EGF2. Consistent with previous results, EGF3 primarily affected dorsal guidance.

### A deletion in the Netrin C-terminal domain resembled a strong *unc-6* loss-of-function

*unc-6(zd144)* was a 132-bp deletion in a region encoding the Netrin C-terminal domain (NTD) (Figure 1 and Table 1). Previous studies indicate that the NCT domain is dispensable for D-V axon pathfinding and instead prevents ectopic axon branching (WANG AND WADSWORTH 2002). *unc-6(zd144)* displayed strong defects in dorsal VD/DD dorsal axon pathfinding and AVM ventral axon pathfinding, although significantly weaker than the *unc-6(ev400)* null (Table 2). the *zd144* deletion includes the exon 11/intron 11 boundary (the third to the last exon) (Figure 1 and Table 1). Possibly, splicing is disrupted in this mutant resulting in a premature stop codon and nonsense-mediated transcript decay, resulting in the *unc-6* loss-of-function phenotype.

### Mutations in predicted N-glycosylation sites in UNC-6

UNC-6 contains three predicted N-glycosylation sites that are conserved in vertebrate Netrin1. The sites are N114 and N128 in the LN domain (Figure 1B and Figure 2D and E), and N423 in the third EGF domain (Figure 1B). We used cas9 genome editing to convert these residues to alanine to eliminate potential glycosylation. *unc-6(lq194)* was N114A, *unc-6(lq196)* was N128A, and *unc-6(lq200)* was N423A. Additionally, the *unc-6(lq194lq197)* was an N114A-N128A double mutant, and *unc-6(lq194lq197lq199)* was the N114A-N128A-N423A triple mutant. All three single mutants and these multiply mutant strains did not display apparent Unc defects. Only the *unc-6(lq194lq197lq199)* triple mutant had significantly more severe VD/DD defects compared to wild type (15.58 versus 15.95 axons reaching reaching the DNC) (Table 2). None of the mutants displayed ventral AVM axon guidance defects (Table 2). Our results reveal that even the loss of all three predicted N-glycosylation sites only had weak effects on dorsal-ventral axon guidance.

### *unc-6 N114*, *N128*, and *N423* mutations interact genetically with *unc-40* and *unc-5*

UNC-40/DCC acts as a receptor with UNC-6 in both dorsal and ventral axon guidance (NORRIS AND LUNDQUIST 2011). Dorsal guidance defects of *unc-40(n324)*, a strong loss of function, were weaker than *unc-6(ev400)*, with an average of 15.76 axons emerging from the ventral cord, and an average of 9.88 reaching the dorsal nerve cord (Table 2 and Figure 3H), similar to what has been previously reported. In double mutants with *unc-40(n324)*, each asparagine-altered mutant significantly enhanced defective growth to the DNC but not the extension of axons emerging from the VNC (Table 2 and Figure 3I).

UNC-5 acts as a receptor with UNC-6 in primarily dorsal axon guidance. In *unc-5* null mutants, very few VD/DD axons emerge from the ventral cord (Table 2) (GUJAR *et al*. 2018). *unc-5(ev480)* is a hypomorphic mutation only affecting the full-length *unc-5A* isoform (MAHADIK AND LUNDQUIST 2023). *unc-5(ev480)* mutants displayed an average of 10.72 VD/DD axons emerging from the ventral nerve cord, and an average of 6.61 reaching the dorsal nerve cord (Table 2). The *unc-6(lq194lq197lq199)* triple mutant significantly enhanced VD/DD defects of *unc-5(ev480)* (Table 2). Each of the single mutants enhanced defects in reaching the dorsal cord (Table 2). Surprisingly, each single mutant and the double mutant significantly suppressed defective emergence from the VNC of *unc-5(ev480)* (Table 2), suggesting a complex genetic interaction with *unc-5*.

*unc-40(n324)* displayed 11% failure of AVM ventral axon guidance (Table 2 and Figure 4C). *unc-6(lq194lq197lq200)* did not significantly enhance AVM ventral axon guidance defects of *unc-40(n324)* (Table 2). The strong loss-of-function *unc-5(e791)* mutants displayed 4% AVM ventral axon guidance defects, whereas the hypomorphic *unc-5(ev480)* displayed no defects (Table 2). The single, double and triple mutants with *unc-5(ev480)* each displayed weak AVM ventral guidance defects, but they did not differ significantly from *unc-5(ev480)* alone. The *unc-6(lq194lq197lq200)* triple mutant did not significantly enhance strong *unc-5(e791)* mutant (Table 2).

N114 and N128 both have carbohydrate linkages in the crystal structure (Figure 5A and B), but N423 does not (Figure 5C). N114 and N128 make no other hydrogen bonds with surrounding residues, but N423 in EGF3 makes three hydrogen bonds (Figure 5). Thus, N423 might weakly perturb the structure of EGF3 in addition to blocking potential N-glycosylation.

**Figure 5.**
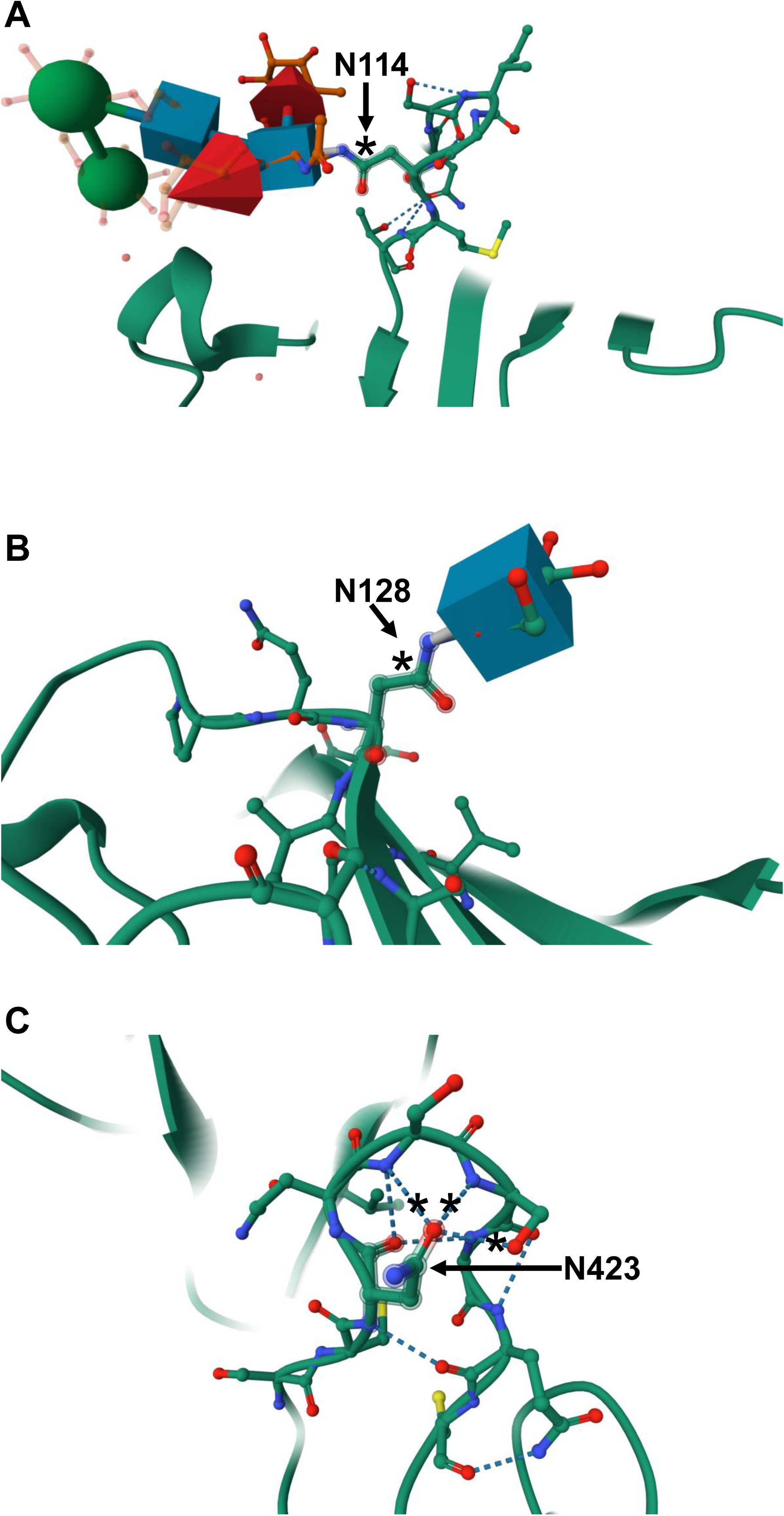
Potential glycosylation sites in UNC-6. See Figure 2 for structure generation. Three conserved potential N-glycosylation sites were found in UNC-6. A) In the UNC-6 crystal structure, N114 has a complex alpha-D-mannopyranose linkage (blue boxes, red cones, and green spheres). B) N128 has a 2-acetomido-2-deoxy-beta-D-glucopyranose linkage (blue box). N114 and N128 do not make other hydrogen bonds. C) N423 has no carbohydrate linkage and instead make three hydrogen bonds with neighboring residues in the EGF3 domain (asterisks).

## Methods

### Genetics

LG I: *unc-40(n324), zdIs5[Pmec-4::gfp]*. LG II: *juIs76[Punc-25::gfp].* LG IV: *unc-5(e791* and *ev480), zdIs4[Pmec-4;;gfp]*. LG X: *unc-6(ev400, e78, zd6, zd17, zd21, zd59, zd61, zd81, zd116, zd121, zd123, zd131, zd144, zd146, zd154, zd156, lq194, lq196, lq200, lq194lq197, and lq194lq197lq199;* LGIV: *zdIs3[Pglr-1::gfp];* LGV: *oyIs14[Psra-6::gfp] V. lqEx1241[unc-6(+)]* is an extrachromosomal array.

To identify axon guidance mutants, we treated animals containing neuron-specific GFP reporters (*oyIs14[Psra-6::gfp] V, zdIs3[Pglr-1::gfp] IV*) with the mutagen EMS (BRENNER 1974) or ENU (DE STASIO AND DORMAN 2001) and then screened their F2 progeny using fluorescence microscopy for animals with altered axon trajectories. Eleven *unc-6* alleles were recovered in screens using *oyIs14[Psra-6::gfp]* following EMS (*zd6, zd17, zd81, zd116, zd121*) and ENU (*zd123, zd131, zd144, zd146, zd154, zd156*) treatment. Three *unc-6* alleles were isolated in screens using *zdIs3[glr-1::gfp]* following EMS (*zd21, zd59, zd61*) treatment. The fourteen new *unc-6* alleles were tested for complementation of the null *unc-6(ev400)* allele using the Uncoordinated phenotype. *lqEx1241[Punc-6::unc-6(+]* is a transgene that rescues the *unc-6* Uncoordinated phenotype and axon guidance defects (HOOPER *et al*. 2025). The *juIs76[Punc-25::gfp]* transgene (JIN *et al*. 1999) was used to visualize the VD and DD GABA-ergic motorneurons (HOOPER *et al*. 2025). *unc-6(ev400) X; juIs76 II; lqEx1241* males were mated to each of the fourteen *unc-6* alleles. Cross progeny containing the *juIs76* transgene (*e.g. zd6/ev400 trans-*heterozygotes) were scored for the Uncoordinated phenotype. Those cross progeny that inherited the *lqEx1241* transgene will be non-Unc even in the case of non-complementation because of the rescuing transgene. Animals that did not inherit the *lqEx1241* transgene would not be rescued and would be Uncoordinated if the new allele failed to complement *unc-6(ev400)*. Both male and hermaphrodite cross progeny with *juIs76* were scored. For each of the fourteen alleles, Unc cross-progeny were detected, indicating failure to complement *unc-6(ev400)* for Unc.

To confirm that the mutations are alleles of *unc-6*, each of the new mutations was crossed into the *lqEx1241 unc-6(+)* rescuing transgene. *lqEx1241* males were mated to each of the fourteen new mutations. In each case, non-Unc male cross progeny were found that were hemizygous for the new *unc-6* mutant and carried the *lqEx1241* rescuing transgene. These males were mated back into the new *unc-6* mutant and non-Unc cross progeny hermaphrodites were selected, which were homozygous for the new *unc-6* allele and harbored the *lqEx1241* rescuing transgene. These were used to create stable, homozygous rescued lines for each of the fourteen new *unc-6* mutants.

### *unc-6* allele sequencing

The *unc-6* locus from each of the fourteen new *unc-6* alleles was amplified by PCR in two overlapping fragments (see Supplemental Materials for primer sequences). The amplified fragments were subject to Oxford Nanopore DNA sequencing (Plasmidsaurus, USA). FASTQ Reads were aligned to the genome using BWA-MEM2 (LI AND DURBIN 2009). The resulting BAM files analyzed on the Integrative Genomics Viewer (ROBINSON *et al*. 2011; THORVALDSDOTTIR *et al*. 2012). Sequence alterations in each of the fourteen new *unc-6* strains were identified (Table 1). Primer sequences used for amplicon sequencing are in Supplemental File 1.

### Scoring dorsal-ventral axon guidance defects

The *juIs76[Punc-25::gfp]* (JIN et al. 1999) transgene was used to score VD/DD GABA-ergic dorsal axon guidance (HOOPER *et al*. 2025). The 19 VD and DD axons grow from the ventral nerve cord (VNC) to the dorsal nerve cord (DNC) to form commissures. In wild type, an average of 16 commissures is observed due to some axons extending in the same commissural fascicle (Table 1 and Figure 3A). For each genotype, the number of axons emanating from the VNC and the number of commissures reaching the DNC was counted (Table 1). 100 animals were scored. One-way ANOVA with Tukey’s post-hoc test was used to statistically compare genotypes and to correct for multiple testing (see Supplemental Materials for all raw data and analysis). ANOVA was used on three groups: *wild type* and *unc-6* mutants; *wild-type, unc-40(n324),* the double mutants with *unc-40(n324)*; and *wild-type, unc-5* mutants, double mutants with *unc-5(ev480)*.

The *zdIs4* and *zdIs5 Pmec-4::gfp* transgenes (ROYAL *et al*. 2005; HOOPER *et al*. 2025) were used to score ventral AVM axon guidance. In wild type, the AVM axon emanates from the cell body directly to the VNC, and then extends anteriorly in the VNC (Figure 4). For each genotype, the percentage of AVM axons failing to emanate directly to the ventral from the cell body was determined. In most cases the defective axons extended anteriorly along the lateral region (Figure 4). 100 AVM axons were scored. In *unc-40,* AVM was sometimes displaced posteriorly due to Q neuroblast and descendant migration defects (HONIGBERG AND KENYON 2000; SUNDARARAJAN AND LUNDQUIST 2012). Only AVMs in the *wild-type* position were scored in *unc-40* mutants. Fisher’s Exact test was used for pairwise statistical comparisons between genotypes. Data and analysis can be found in Supplemental File 2.

### Cas9 genome editing

The potential glycosylation sites N114, N128, and N423 in *unc-6* were changed to alanine residues using Cas9 genome editing and standard techniques (DICKINSON *et al*. 2013; NORRIS *et al*. 2015). See Supplemental Material for the sequences of synthetic guide RNA *unc-6* inserts, the single-stranded oligonucleotide repair templates, and sequences of the genome-edited *unc-6* mutants. A mix of sgRNAs, a single stranded oligonucleotide repair template, and Cas9 enzyme was injected into the gonads of N2 animals, along with the *dpy-10(cn64)* co-CRISPR reagents (EL MOURIDI *et al*. 2017). Genome-edited animals were identified using PCR with primers specific to the recoded regions introduced from the repair template. The genome edits were confirmed by PCR and sequencing. Genome editing reagents were produced by InVivo Biosystems (Eugene, OR, USA).

Individual *lq194 N114A, lq196 N128A,* and *lq200 N423A* were generated in a *wild-type* N2 background. *lq197 N128A* was induced in the *lq194 N114A* background to create the *lq194lq197* double mutant. This required the use of a novel repair template oligonucleotide that corresponded to the recoded sequence of the N114A genome edit. *lq199 N423A* was induced in the *lq194lq197* background to create the *lq194lq197lq199* triple mutant. Primer sequences for genome editing can be found in Supplemental File 1.

## Discussion

### Mutations in the Laminin N-terminal domain

Four of the new *unc-6* mutations are missense mutations that alter distinct domains of UNC-6. Previously-described mutations in the *unc-6* LN domains perturbed both dorsal and ventral growth (LIM AND WADSWORTH 2002; XU *et al*. 2009). *zd17* (S120F) and *zd146* (G289D) in the LN domain had distinct effects on dorsal-ventral axon pathfinding. VD/DD dorsal guidance defects and AVM ventral guidance defects in *unc-6(zd146)* were comparable to the null *unc-6(ev400)*, indicating that the conserved glycine 289 is needed for all UNC-6 functions. The importance of this residue is highlighted by mutation of the residue in Laminin α2 causing limb-girdle-type dystrophy in humans (GAVASSINI *et al*. 2011) and in Laminin β1 causing cardiac development defects in *Drosophila* (HOLLFELDER *et al*. 2014). By contrast, *unc-6(zd17)* was a hypomorphic *unc-6* mutation that more strongly disrupted ventral axon pathfinding than dorsal. S120 makes hydrogen bonds with an adjacent beta strand in a domain outside of the “jellyroll domain” that is glycosylated at N114 (Figure 2E). However, mutation of N114 did not disrupt dorsal or ventral guidance, arguing that another aspect of the S120F mutation is perturbing ventral guidance. A distinct region of the LN domain of Netrin-1 is required for interactions with the 4^th^ fibronectin domain of DCC and Neogenin (XU *et al*. 2014), potentially explaining the role of the LN domain in both dorsal and ventral guidance. The S120F mutation in *zd17* might specifically disrupt interactions with UNC-40 or another receptor affecting ventral but not dorsal guidance. LN domains also interact with one another, forming the Laminin lattice network in the basal lamina (SHAW *et al*. 2021; KULCZYK *et al*. 2023). *zd17* might affect interaction of UNC-6 with Laminin, which might be required for ventral guidance.

### Mutations of the three EGF domains

EGF3 is specifically involved in dorsal axon guidance (LIM AND WADSWORTH 2002; NORRIS AND LUNDQUIST 2011). We found that *unc-6(e78),* which alters a disulfide-bond-forming cysteine residue in EGF3 (Figure 2C), had strong defects in dorsal VD/DD axon guidance but did not strongly affect ventral AVM growth (Table 2), consistent with these results. EGF3 of Netrin-1 interacts physically with the 5^th^ fibronectin domain of DCC (XU *et al*. 2014). In *C. elegans*, UNC-40 is involved in both dorsal and ventral growth (NORRIS AND LUNDQUIST 2011; NORRIS *et al*. 2014), so it is possible that potential EGF3 interactions with UNC-40 are specific for dorsal growth.

EGF2 is required for both dorsal and ventral growth (LIM AND WADSWORTH 2002). *unc-6(zd116)* altered a disulfide-forming cysteine in EGF2 (Figure 2B). *unc-6(zd116)* mutants had defects in dorsal VD/DD and ventral AVM axon guidance that were significantly less penetrant than the *unc-6(ev400)* null. Our data indicate that EGF2 is required for both dorsal and ventral guidance, consistent with previous structure-function studies.

*unc-6(zd154)* affected a disulfide-forming cysteine in EGF1 (Figure 2A), and mutants had strong defects in dorsal VD/DD axon guidance similar to the *unc-6(ev400)* null. Effects on ventral AVM axon guidance were weaker than *unc-6(ev400)*. Our data show that EGF2 is involved in both dorsal and ventral guidance, but with a more pronounced role in dorsal axon guidance. This observation is consistent with Alphafold predictions of the UNC-5 2^nd^ immunoglobulin domain interacting with EGF1 (PRIEST *et al*. 2024) possibly to mediate dorsal growth away from UNC-6.

In sum, our analysis of domain-specific missense mutations suggests that the LN domain, EGF1, and EGF2 are required for both dorsal VD/DD and ventral AVM axon guidance to varying extents, and that EGF3 is required for dorsal VD/DD axon guidance with a minimal role in ventral guidance (Figure 6). The S120F region in the LN domain might predominantly be required for ventral AVM guidance. One caveat is that domain-specific mutations might have secondary effects on UNC-6 overall structure and stability, making it difficult to assign specific functions to specific domains. However, the previous structure-function studies (LIM AND WADSWORTH 2002; XU *et al*. 2009) and physical interactions (XU *et al*. 2014; PRIEST *et al*. 2024) are consistent with domain-specific functions.

**Figure 6.**
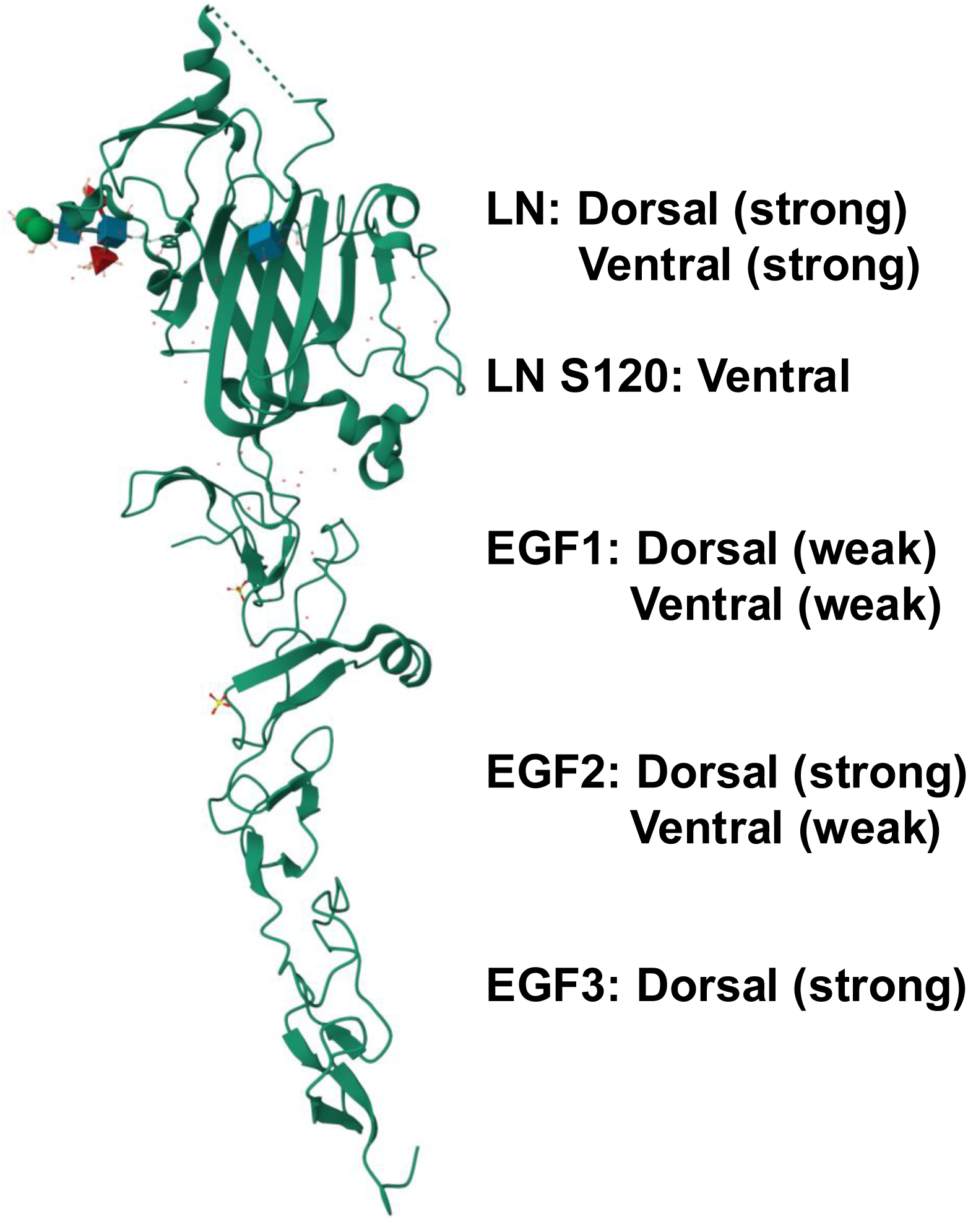
Roles of UNC-6 regions in dorsal VD/DD and ventral AVM axon guidance. See Figure 2 for structure generation. Disruption of the LN domain strongly perturbed both dorsal VD/DD and AVM ventral guidance, and alteration of the S120 residue in the LN domain predominantly perturbed ventral AVM guidance. Mutation of the EGF1 domain perturbed both dorsal and ventral guidance but with weaker effects on ventral guidance. Disruption of the EGF2 domain weakly perturbed both dorsal and ventral guidance. Mutation of EGF3 predominantly disrupted dorsal VD/DD guidance.

### Potential N-glycosylation site mutations interacted genetically with *unc-40* and unc-5

UNC-6 N114, N128, and N423 are predicted N-glycosylation sites. Indeed, N114 and N128 had carbohydrate linkages in a crystal structure whereas N423 did not (PRIEST *et al*. 2024) (Figure 2). Mutations altering each individual site to alanine had no detectable effect of dorsal-ventral axon guidance, nor did double or triple mutants. However, N114, N128, and N114-N128 enhanced VD/DD dorsal axon guidance defects of *unc-40* mutants. Possibly, these sites function in parallel to UNC-40 in dorsal axon guidance.

Surprisingly, the single mutants and the N114A-N128A double mutant suppressed the hypomorphic *unc-5(ev480)* mutation for the number of axons emanating from the VNC. This observation might reflect the balance of UNC-5 and UNC-40 response to UNC-6 in dorsal VD/DD outgrowth in the Polarity-Protrusion model (NORRIS AND LUNDQUIST 2011; GUJAR *et al*. 2018). In the absence of UNC-5 repulsive signaling, UNC-6 might draw the axons back to the VNC via attractive signaling and UNC-40 (NORRIS AND LUNDQUIST 2011; NORRIS *et al*. 2014; GUJAR *et al*. 2017; GUJAR *et al*. 2018; GUJAR *et al*. 2019; MAHADIK AND LUNDQUIST 2023; HOOPER *et al*. 2025). This explains why the dorsal VD/DD defects of *unc-5* mutants are stronger than *unc-6* and are suppressed by *unc-40* (NORRIS AND LUNDQUIST 2011). Possibly, UNC-6 function is slightly perturbed in N114A, N128A, and N423A, resulting in reduced UNC-5 signaling and less growth back toward the VNC. *unc-5(ev480)* with the triple mutant N114A-N128A-N423 showed significantly fewer VD/DD axons emerging from the VNC, indicating that the cumulative effect of the three mutants begin to have more severe loss of *unc-6* function that enhances *unc-5(ev480)*. The number of VD/DD axons reaching the VNC in *unc-5(ev480)* was significantly reduced by each single mutant and the double and triple mutant. Possibly, once the axons have left the VNC, the mutations UNC-6 acts with UNC-5 and the N114A, N128A, and N423A mutations enhance the phenotype of the *unc-5(ev480)* hypomorph.

For AVM ventral guidance, the triple N114A-N128A-N423A mutant did not significantly enhance *unc-40(n324)* (Table 2). However, the mutations did cause AVM ventral guidance in the context of the *unc-5(ev480)* hypomorph to an extent similar to the *unc-5(e791)* null. For example, *unc-5(ev480)* showed no AVM ventral guidance defects whereas the *unc-5(ev480); N114AN128An423A* mutants showed 7%, similar to *unc-5(e791)*. Together, these data indicate that N114, N128A, and N423 slightly perturb UNC-6 function revealed in the context of genetic interactions with *unc-40* and *unc-5*. Biochemical and structural analyses reveal that UNC-5 required heparin for strong binding to UNC-6, which also excludes UNC-40 interaction (PRIEST *et al*. 2024). The relation of this result to N114A and N128A is unclear, but it is possible that glycosylation of UNC-6 is required for robust interaction with UNC-5. It is also possible that these mutations each slightly perturb the structure or stability of UNC-6 in a cumulative fashion. In any case, it is unlikely that N423 is a *bona fide* glycosylation site and instead might be involved in hydrogen bonding in the EGF3 domain structure (PRIEST *et al*. 2024) (Figure 5C).

In sum, these studies delineate the roles of specific domains of UNC-6/Netrin in dorsal VD/DD axon guidance and ventral AVM axon guidance using missense mutations in the endogenous *unc-6* locus. The LN domain was required for both dorsal and ventral guidance, but S120 in the LN domain was specific to ventral AVM axon guidance. EGF1 and EGF2 were required for both dorsal and ventral guidance, and EGF3 was specific to dorsal guidance as previously reported. Finally, N-linked glycosylation at N114 and N128 was not required for UNC-6 function in a *wild type* background, but modified dorsal and ventral axon guidance in *unc-40* and *unc-5* mutant backgrounds. Our results will inform future studies of how these distinct domains interact with the guidance receptors UNC-40, UNC-5, and other extracellular molecules to control dorsal-ventral axon guidance.

## Acknowledgements

We thank C. Chiu, B. Kahoussi, B. Lee, K. Sabater and T. Smith for undertaking the screens that identified the *unc-6* alleles. Some strains were provided by the CGC, which is funded by NIH Office of Research Infrastructure Programs (P40 OD010440).

