## Supplemental File 1 for "Domain-specific mutations in *unc-6/Netrin* differentially affect dorsal-ventral axon pathfinding in *Caenorhabditis elegans*"

Primer sequences used in this work

| Name | Purpose | Sequence |
| --- | --- | --- |
| unc-6amp1F | Amplicon sequencing of *unc-6* alleles | agtcaatttccagattgtgtc |
| unc-6amp1R | Amplicon sequencing of *unc-6* alleles | gtgctgactgaccagtaggta |
| unc-6amp2F | Amplicon sequencing of *unc-6* alleles | ggtggtagccattcttccgtg |
| unc-6amp2R | Amplicon sequencing of *unc-6* alleles | gaacagcttgatatggctgcc |
| N114AsgRNA | sgRNA insert for the N114A genome edit | cgattggaaacatgacatgctgg |
| N128AsgRNA | sgRNA insert for the N128A genome edit | ccaagtttgagcccacaaaacgt |
| N423AsgRNA | sgRNA insert for the N423A genome edit | aaaagctgcaaccaatcatcggg |
| N114AssODN | Repair template oligonucleotide for the N114A genome edit | gtgatgctagaaaccatttccaatcc  catccagcctccctccttaccgacctt  aattctatcggagctatgacttgttgg  gtctctactccaagtttgagcccacaa  aacgtttcactcactttg |
| N128AssODN | Repair template oligonucleotide for the N128A genome edit | ttcgattggaaacatgacatgctgggttt  ccactccgtccctttctccacaggccgtc  tcccttaccctttcccttggaaagaaattc  gaacttacctacgtttccatgcatttttgct  ctagactcccagattcaatggcactttac  aagtctgctgac |
| N114N128AssODN | Repair template oligonucleotide for the N128A genome edit in the N114A background | ttctatcggagctatgacttgttgggtc  tctactccgtccctttctccacaggcc  gtctcccttaccctttcccttggaaag  aaattcgaacttacctacgtttccatg  catttttgctctagactcccagattcaa  tggcactttacaagtctgctgact |
| N423AssODN | Repair template oligonucleotide for the N423A genome edit | tcaaaaccacttaagcttgtggatgtc  atccagtcggtagcctcggtaagtctt  gtgctcagtcatcgggtcagtgcgtct  gcaagcctggagtcactg |
